# Posterior parietal cortex maps progress along routes sharing the same meta-structure

**DOI:** 10.1101/2024.06.17.599318

**Authors:** Alexander B. Johnson, Douglas A. Nitz

## Abstract

Neurons of posterior parietal cortex were recorded as rats performed a working memory task within a network of intersecting paths. The specific routes utilized in task performance provided opportunity to contrast responses of posterior parietal cortex sub-populations to linear and angular velocity with more complex responses that map route progress. We found evidence for the presence of posterior parietal cortex neurons that generalize in their firing patterns across routes having the same shape but opposite action series. The results indicate that posterior parietal cortex has the capacity to generalize the mapping of route progress independent of the specific actions taken to move through those routes. We suggest that such encoding can form the basis for learning the meta-structural organization of a non-random path network structure, such as that commonly found in cities.

## Introduction

For many environments, travel is constrained and/or optimized by moving along pathways. When multiple pathways are connected to one another, the intersections and orientations across multiple pathways form what can be considered as a ‘path network’, and its structure can range from highly regular to nearly random. Efficient movement within environments containing a path network can be facilitated through prior knowledge of the path network’s structure. For environments that have some degree of repeating geometric structure, multiple different routes may have the same shape despite being defined as connecting different locations or being oriented differently from one another. In principle, memories for the specific shape of a route can form the basis for learning the organization of a path network structure when a view of the full network cannot be taken. In this conceptualization, the shapes, start and end locations, and orientations of routes can be thought of as puzzle pieces which fit together to form the picture of the path network.

Clues to how this mental process manifests in the nervous system come from humans who have suffered brain damage and exhibit topographic amnesia/agnosia. This syndrome is characterized by a profound inability to assess one’s location with respect to landmarks in the environment, the inability to confidently navigate previously learned routes, and the inability to successfully learn new routes (De Renzi et al., 1977). These people often have the locus of their brain damage in or near the posterior parietal cortices (PPC) (Takahashi et al., 1997; Takahashi and Kawamura, 2002).

In rats, neurons of PPC are known to encode progress along a route. The frame of reference which these neurons respect is that of a route as having a particular shape irrespective of its environmental location and orientation, independent of the scenes associated with its traversal, and independent of the series of navigational actions taken to complete it (Nitz, 2006; Whitlock et al., 2012). Some work suggests that neural representations for progress along a path are scalable such that the same path shape taken over different total distances will yield the same series of ensemble activity patterns in PPC (Nitz, 2006; Nitz, 2012). Notably, PPC ensembles discriminate route locations sharing the same locomotor action such as left or right turns (Nitz, 2009), and PPC route progress encoding develops over just a few traversals of that route (Nitz, 2006). Other findings indicate that PPC neuron activity can also be explained through their tuning to different linear and angular velocities integrated across differential offsets in time (Alexander et al., 2022). Integration of this form can, in principle, form the basis for learning route shape through locomotor experience alone. The patterns of spatial tuning as described in PPC can be contrasted with the trajectory-specific encoding of specific environmental locations observed for “place cells” of sub-region CA1 of hippocampus (Frank and Wilson, 2001; Wood et al., 2001; Ferbinteanu and Shapiro, 2003).

The known activity patterns in PPC form a partial explanation for experience-based learning of a path network structure (Chrastil and Warren, 2015). It is possible, though not yet observed, that shared meta-structural features of routes, such as their shape, might facilitate learning of a path network structure as well. Shared meta-structural features of a path network could preserve the shape of a route while the action sequence performed differs. In a simplified case of a squared grid environment, any two complementary action sequences (e.g. four consecutive 90-degree lefts or rights) generate the same shape, a rectangle, but are mirrored in orientation with respect to deviations from the starting orientation. Such route shapes, reflecting the navigational affordance structure of the environment, will recur from a multitude of starting locations. Learning that the same shape recurs in the environment evidences an understanding of the larger environment topography, such as a squared grid pattern, and the navigational predictions that accompany that. While PPC neuron ensembles can encode progress along routes of a particular shape as defined by an action sequence, it is presently unclear to what extent the encoding of route progress more generally translates onto routes that differ in their action sequence but are the same in their shape. This is the key question addressed in the present work.

To examine PPC neuron mapping of actions versus route locations, we analyzed 236 identified single-units recorded from PPC as rats navigated an interconnected path-network, the ‘Triple-T’ maze. Performance of a working memory navigational task within the environment allowed us to compare PPC responses to the linear and angular velocity dynamics associated with turning actions with PPC responses to analogous locations across routes that were identical in shape while differing in their constitutive action sequence. We observed a sub-population of PPC neurons exhibiting opposite firing patterns for pairs of routes that demand opposite action sequences. A second sub-population mapped progress through pairs of routes sharing the same shape, despite their differences in the locomotor action sequences they demanded. Together, the findings extend prior work on spatial tuning of PPC neurons in freely-moving animals to reveal spatial tuning reflective of meta-structural similarities among pathways that differ with respect to actions, headings and environmental locations. We suggest that this form of tuning could form the basis for the learning of environmental path-network structure in the absence of a top-down, map-like view. These spatial representations furthermore suggest a neural basis for the conceptualization of structural similarities across unique spaces.

## Methods

### Rats

Subjects were 5 male Sprague Dawley rats. From these rats, 236 PPC neurons were recorded and isolated. Animals were all between 6 and 10 months old at the beginning of training and were singly housed in standard plastic cages. The vivarium was kept on a 12-hr light-dark cycle. Rats were initially on an ad libitum feeding schedule, however after initial exposure to the recording room animals were food restricted to maintain a weight between 85% and 95% baseline to maintain motivation throughout the task. Experimental protocols followed all AALAC guidelines and were approved by the Institutional Animal Care and Use Committee guidelines at the University of California, San Diego.

### Surgery

Following one month of pre-surgery training on the triple-T working memory task. Rats were surgically implanted with custom built microdrives each equipped with bundles of 12.5 micron tungsten wires spun in groups of 4 into tetrodes. Rats were implanted unilaterally or bilaterally with microdrives positioned dorsal to PPC with wires initially positioned approximately 0.5mm deep into cortex. Coordinates for implants were determined through referencing the Paxinos and Watson Rat Brain Atlas (Paxinos and Watson 2014). PPC coordinates relative to bregma were centered at A/P -3.8 mm, M/L ±2.3 mm, D/V 0mm - 0.5mm. The microdrive implant allowed wires to be moved ventrally through PPC across days in 40um increments. Tetrode locations were verified post-hoc using Nissl-stained tissue by the presence of visible tracts in the tissue (Supplemental Figure 2). All surgeries were performed in compliance with the Institutional Animal Care and Use Committee guidelines at the University of California, San Diego.

### Triple-T Maze Environment

Experiments were conducted on a “triple-T” path-network maze. The track (Figure 1A; 8-cm-wide pathways, overall perimeter 1.6 m × 1.25 m in length and width, painted black) stood 20cm high in the middle of the recording room. The track edges were 2 cm in height, allowing an unobstructed view of the environment’s boundaries and associated distal visual cues. Access to certain areas of the maze was restricted by placing painted black cans at key junctions. The placement of these and the locations of food reward sites organized navigational behavior into 4 internal pathways each measuring 140 cm in length with forced left/right turn decisions located 51 cm, 87 cm, and 118 cm along each pathway (Figure 1B, C). Paths 1 and 4 and paths 2 and 3 demand opposite turn sequences and were mirror images of each other. Two perimeter “return” routes formed a third set of mirror-image route shapes and flanked the internal portions of the maze. Each was 197 cm in length and demanded two left or two right turns.

**Figure 1.**
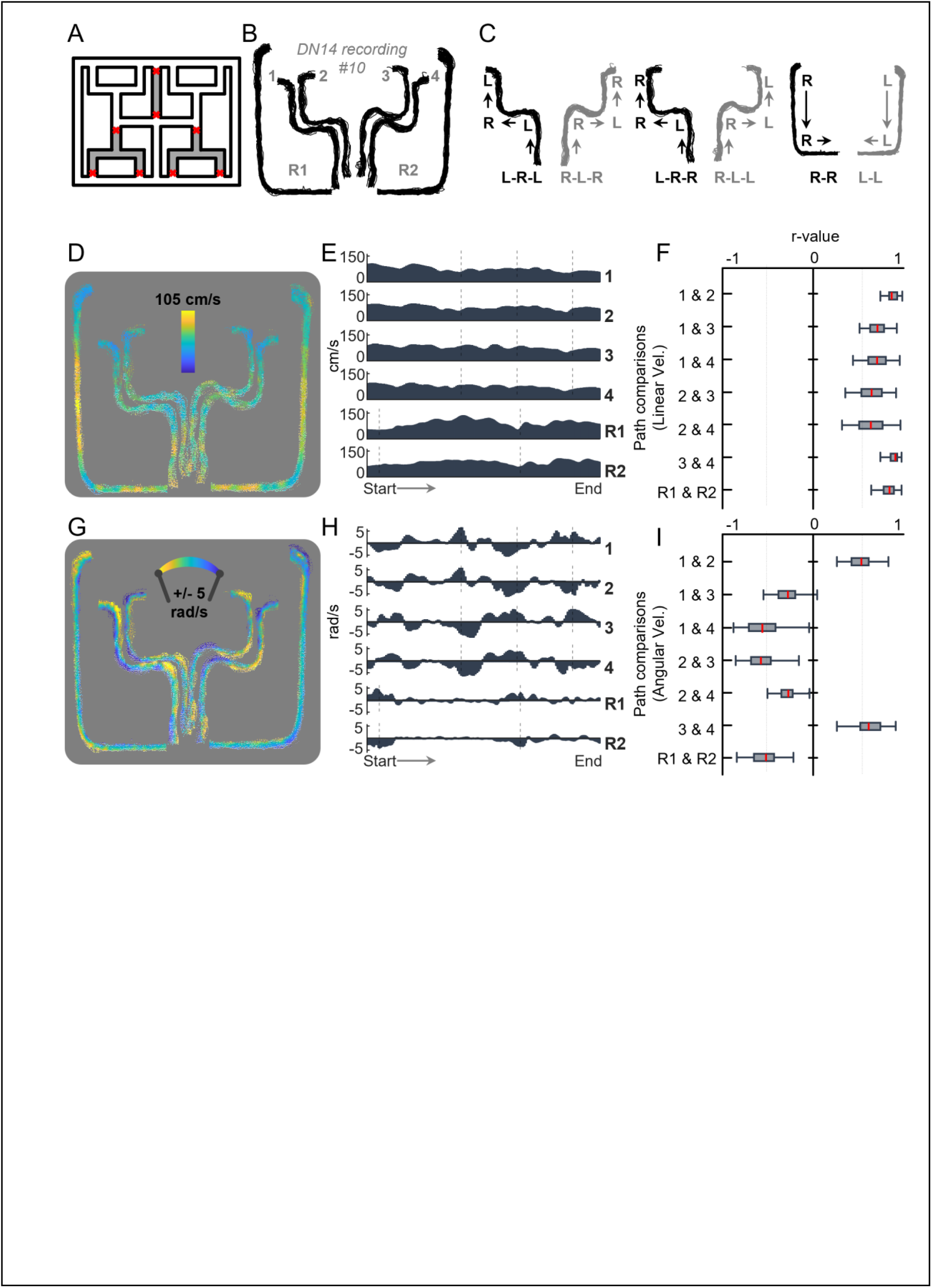
Components of Self-Motion During Spatial Working Memory Task. A) Schematic of triple-T environment configured to the find-all-4 task. Blockers, marked by red X’s, were implemented to define the accessible portion of the environment. B) Tracking data collected from one example recording session are shown with overlapping portions of traversals separated to illustrate the labelling of individual uninterrupted runs based on their terminal location. C) Three path-path pairs utilize opposite action sequences. Individual internal pathway pairs 1 and 4, pathway pairs 2 and 3, and the external pathway pair of R1 and R2 can be considered to be mirror images of each other despite the utilization of opposite turning sequences. D) Individual traversals from an example recording are illustrated and color-coded with respect to the animal’s measured linear velocity. E) Linearized velocity-vectors for each of the routes traversed illustrated in D averaged across all traversals for that recording. F) Pairwise correlation values for each recording’s averaged linearized velocity vectors. Demonstrating strong similarity in measured velocities across all pathway pairs. G-I) Same as D-F, but for measured angular velocity. Demonstrating strong dissimilarity in measured velocities for path-pairs 1&4, 2&3, and R1&R2.

### Spatial Working Memory Behavior Task

Rats were habituated to the “triple-T” maze for 2 periods of about 30 minutes prior to training. During the first habituation period the animal had access to the entire maze without any blockers or other obstructions present. The second habituation period took place the following day and utilized blockers to restrict movement to the corridors used in the task (Fig 1A). Following habituation rats were trained to traverse one of the four available internal pathways for food-reward. Following the collection of the food reward animals learned to utilize the perimeter routes of the maze to return to the ‘main stem’, the shared portion of each internal route, and begin another traversal for another food reward. Rats were permitted to choose whichever route back to the ‘main stem’ they preferred and were permitted to turn around only on the perimeter pathways. Rats often did not change their direction however, often restricting their behavior to a single direction for each position of the maze and maintaining consistent self-motion (Figure 1 D-I). Once animals regularly performed 80% or more non-interrupted traversals of all four internal pathways a reward schedule was implemented. The reward schedule started with each of the four possible internal routes being baited with a food reward at the end location. The rat had to gather all the available food rewards before progressing to the next ‘block’ of trials where the food rewards were replaced. Each block required the rat to obtain each of the 4 potential rewards in any order and permitted for ‘error’ trials where no reward was obtained, however, rats quickly learned this find-all-4 rule and performed the task reliably quickly and with high accuracy (Supplemental Figure 1).

### Recording Sessions

Each microdrive implant had one electrical interface board (EIB-16 Neuralynx) connected to an amplifying headstage (20X, Triangle Biosystems). Raw signals were initially amplified (50X), and high-pass filtered (>150Hz) and brought into a dedicated recording computer running Plexon SortClient software. Here, the signal was digitized at 40kHz, band-pass filtered (0.45 – 9kHz), and amplified between 1X and 15X to fit the shape of detected waveforms (for a total of 1,000X – 15,000X). Over time tetrode wires were moved in 40um steps ventrally through brain tissue to maximize the number of unique neural units recorded across days from each animal. Single units were identified and isolated by hand using Plexon OfflineSorter software. Key waveform parameters for separation were peak height, peak-valley distance, energy, average voltage, and principal components.

Animal position data was collected at 60Hz using a ceiling-mounted camera, mounted 305cm above the recording room floor. Colored LED lights affixed to the implants of recorded animals were tracked using Plexon CinePlex Studio software to obtain X, Y coordinates using customized MATLAB software. Lights were approximately 4.5cm apart and were positioned perpendicular to the heading of the animal. Recordings lasted for approximately 30 minutes each during which time animals performed on average 20 blocks of trials, however the total number of trials varied by day. Recording sessions where rats performed under 6 blocks of trials were not included in the final dataset.

### Histology

Rats were perfused intracardially with a solution of 4% w/v paraformaldehyde in PBS during deep anesthesia. Brains were removed and sectioned into 30um slices. Brain slices were Nissl-stained to identify the location, trajectory, and depth of tetrode wires in PPC. Boundaries of PPC were defined based on previous electrophysiological studies and in accordance with Paxinos and Watson atlas (Paxinos and Watson, 2014). Electrodes were determined to be in PPC at the time of recording from post-hoc verification where the terminus location of each tetrode was identified and recorded wire-movement across days was used to calculate the anatomical position for each recording. All tetrodes were determined to have been in the PPC at the time of recording for the units to be included in this study.

### Identification of Clean Traversals

To identify traversals made on the triple-T maze that demonstrated clean and uninterrupted running, custom MATLAB graphical interfaces were utilized. The user defines, in space, the starting and ending ‘gates’ for each route defined for analyses. From those gates, the MATLAB script automatically extracts traversals with sustained running speed at or above 3cm/s between them. The user then verifies each individual run to ensure there are no deviations from uninterrupted stereotyped running behavior. The selection of clean runs results in the data presented in Figure 1B. Trials were dropped from analysis for stalling, or non-stereotyped behavior (e.g. turning around). The defined routes were the four internal routes toward potential reward sites or perimeter routes leading to the internal entrance (Figure 1A).

### Route and Space Referenced Positional Rate Vectors

To analyze the action and spatial correlates for each neuron, individual neuron activity was mapped onto the position of each route using custom MATLAB scripts. Uninterrupted traversals were used and fitted to a template with ∼1cm spatial bins extending from the start of each route to the end. Firing rates were then calculated for each bin by dividing the total number of spikes by occupation time. Activity patterns were then smoothed with a Gaussian filter (σ = 6 cm AUC = 1).

Like the linearized route-referenced positional rate vectors, two-dimensional firing ratemaps were constructed for each neuron for the entire space of the maze for the entire experiment. Tracking samples associated with a velocity at or above 3cm/s the X, Y coordinates are identified and the number of spikes and occupancies for each spatial bin were determined. The mapping of spike counts is divided by the mapping of occupancies and multiplied by 60 to yield an estimate of spikes/second for each spatial bin. This process is done for the entire experiment and averaged across each identified X, Y position. Raw two-dimensional ratemaps were smoothed with a gaussian filter (σ = 6 cm2 AOC = 1).

### Generalized Linear Model

A series of GLMs were computed to assess the impact self-motion had on the activity profiles of individual neurons as rats performed the triple-T task. Linear and angular velocity were chosen as predictors of firing rates as these self-motion measures are often associated with PPC firing rates (e.g., Alexander et al. 2022; Whitlock et al., 2012). A separate “complete” GLM cGLM using linear and angular velocity as predictors was constructed to model each neuron’s linearized max-normalized positional firing rate vector for each internal and external route (using glmfit.m in Matlab). Coefficients were calculated for both linear and angular velocities (glmval function in MATLAB) to reconstruct the activity profile which was used to calculate the fit between the actual firing rate vector and the cGLM output assessed using the normalized mean squared error (‘NMSE’, as an output from the function ‘goodnessOfFit’ in MATLAB). Both predictors had their relative contribution in modelling tested using the accuracy of the GLM as a metric. To accomplish this, a partial GLM (pGLM) was fit to each neuron in the same manner as above with the exclusion of either linear or angular velocity from the model. Kruskall-Wallis tests with post hoc Boneferonni corrections were made comparing the distribution of values derived from the pGLMs relative to their respective cGLMs. To compare across groups of identified neurons, NMSE scores derived from pGLMs were compared using a 2-tailed t-test. cGLM NMSE scores were used to normalize the change in model fit for each pGLM, and the t-test was done on the average proportional change of NMSE for the pGLM calculated for each route.

### Correlation Analyses of Linearized Firing Rates

Individual neurons had their activity patterns compared across identical length across the 4 internal routes and for the 2 external routes. Equal-length linearized positional firing rate vectors were used to calculate a Pearson’s r value as a measure for similarity. Reliability in positional firing for each route was calculated by comparing the odd numbered traversals’ positional firing rates to the even numbered traversals’ rates. Second, a comparison across routes was calculated for each route-pair by comparing the mean firing rates from each route to each other route of equal length.

A distribution of correlation values under randomization was also calculated by rotating the firing rate vectors for each traversal (repeated 100 times for each neuron with different random degrees of rotation). This generated a larger sampling of bootstrapped data from which a distribution of correlation values expected by chance was determined. We took the mean plus or minus two standard deviations for each control distribution to denote significance. Neurons that exhibited both above chance reliability within each route, and exhibited inter-route correlations above chance were classified as positive (+) neurons for each route comparison. Neurons that exhibited both above chance reliability within each route, and exhibited inter-route correlations below chance were classified as negative (-) neurons for each route comparison. If neurons did not exhibit the above chance reliability for either route being considered, the neuron was not classified as either (+) or (-) regardless of inter-route correlations.

## Results

### Behavioral Dissociation of Locomotor Action Series from Progress Through Similarly Structured Routes

Five adult male Sprague Dawley rats were trained to perform a spatial working memory task within the ‘triple-T’ path network (Olson et al., 2019; Olson et al., 2017; Figure 1A). The triple-T task entails the successive utilization of four internal pathways (1-4 in Figure 1B) from a common starting point to reach each of four food reward locations. Between internal path traversals, the animal returns from any of the four food reward locations via either of two return pathways along the perimeter (R1, R2). Thus, the task demands spatial working memory for which paths have been visited in sets of four path-traversal blocks.

The animal is free to utilize any possible ordering of the four internal paths for any given block. Delivery of food reward at the end of any internal path is predicated on the animal not having returned to that location along that path prior to visiting each of the four food reward locations. The animal must eventually reach all four food reward locations within a given block. Therefore, ‘perfect’ blocks are composed of exactly four internal path traversals. Blocks with errors are composed of both rewarded and unrewarded path traversals, typically between 5 and 6 in number.

Each internal path is composed of three turn locations with turn 2 being forced (no L versus R option). Distances between turns are the same for all four pathways, yielding structural similarity in shape for specific path pairings. Each return path is composed of two turn locations, both forced. Figure 1C depicts tracking data for uninterrupted runs along each of the four internal and two return pathways, highlighting the fact that paths 1 and 4, paths 2 and 3, and paths R1 and R2 entail movement along mirror-image path shapes demanding opposite series of L versus R turns. Rats become very proficient in this task (Supplemental Figure 1A, B) collecting a reward for about 84% of all traversals (s.d. = 6.88%). Animals utilized several strategies including the use of the shorter return arm to return back to the main stem of the maze (Supplemental Figure 1C mean = 0.9197 s.d. = 0.1075) and alternation at the first L/R turn site common to all four internal paths (Supplemental Figure 1F mean = 0.8973 s.d. = 0.0916). Despite strong performance and the spontaneous development of these navigational strategies, rats tended not to repeat specific internal path sequences (Supplemental Figure 1D).

Animals navigated at consistently high speeds (Figure 1D, E) and with consistent angular velocities (Figure 1G, H). The profiles of linear velocities were seen to be consistent across routes (Figure 1F) whereas the profiles for angular velocities were seen to differ dramatically for path 1 versus 4, path 2 versus 3 and path R1 versus R2 (Figure 1I). In this way, the task dissociates navigational locomotor actions in the form of turns from progress through the spatial extents of similarly structured and equal-length routes.

### Positional Rate Vectors for PPC Neurons Reveal Mapping of Both Action and Route Progress

For each isolated PPC neuron a positional firing rate vector was created for each of the four internal and two return paths based on those path traversals completed without interruption. Position along all four internal paths was organized into 140 spatial bins approximately 3.5cm apart with bins 51, 87, and 118 representing the peaks of the three turns. Similarly, position along the two return paths is organized into 196 bins with bins 15 and 127 representing the peaks of the two turns. This resulting linearization of the spatial firing profile organized the data for subsequent application of the rate vector correlation and generalized linear modeling (GLM) analytical approaches considered below.

For many PPC neurons, positional firing rate vectors reveal apparent tuning to locomotor actions (L or R turns) as has been described in prior publications (McNaughton et al., 1994; Nitz, 2006; Whitlock et al., 2012; Wilber et al., 2014; Goard et al., 2016). Figure 2A depicts rate vectors for three such neurons along all six paths as well as the correlations in rate vectors for all internal path combinations and the R1/R2 return paths combination. For neurons seeming to encode locomotor actions, rate vector correlations are high for path combination 1,2 and path combination 3,4 where distinction in the form of angular velocity (L/R turn type) is seen only at the very end of the paths.

**Figure 2.**
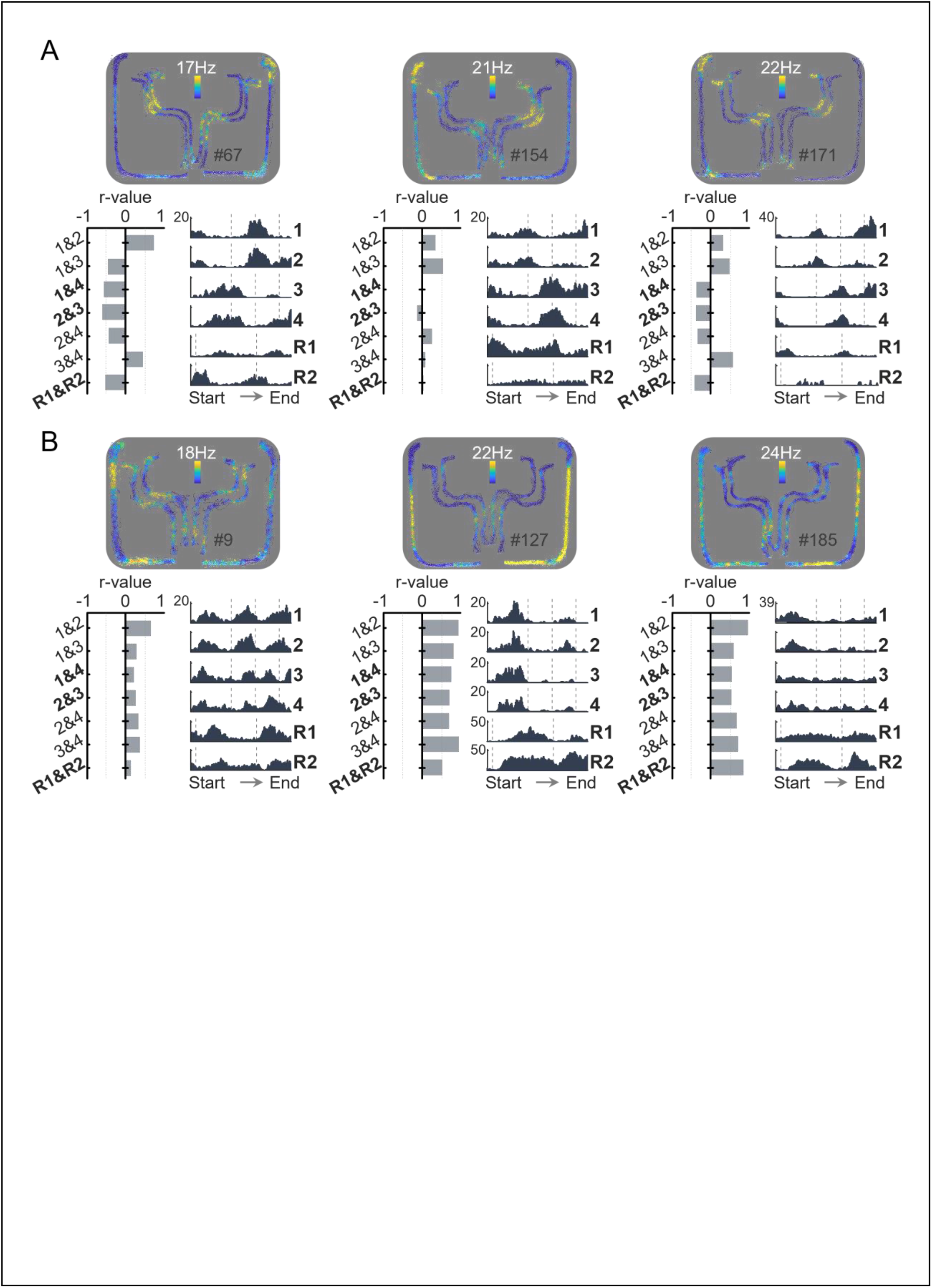
Example Parietal Cortex Neuron Responses on Triple-T Maze. Example neuron responses as recorded on the triple-T presented as 2-D ratemaps with no-firing color-coded as dark blue, and maximal firing as indicated above each example and color-coded as yellow. Below is the mean positional firing rate for each linearized path (right) and the pairwise path-path correlation profile (left). A) Three example neurons demonstrating putative self-motion responses. The leftmost example neuron responds as the animal executes right turns. The middle and rightmost example neurons increase activity as the animal executes left turns. B) Three example neurons highly correlated positional rate vectors across route pairs that demand opposite turn sequences (e.g., 1 vs 4, 2 vs 3, R1 vs R2). Each neuron exhibits similar activity patterns across all internal routes (1-4) and across return routes R1 and R2.

**Figure 3.**
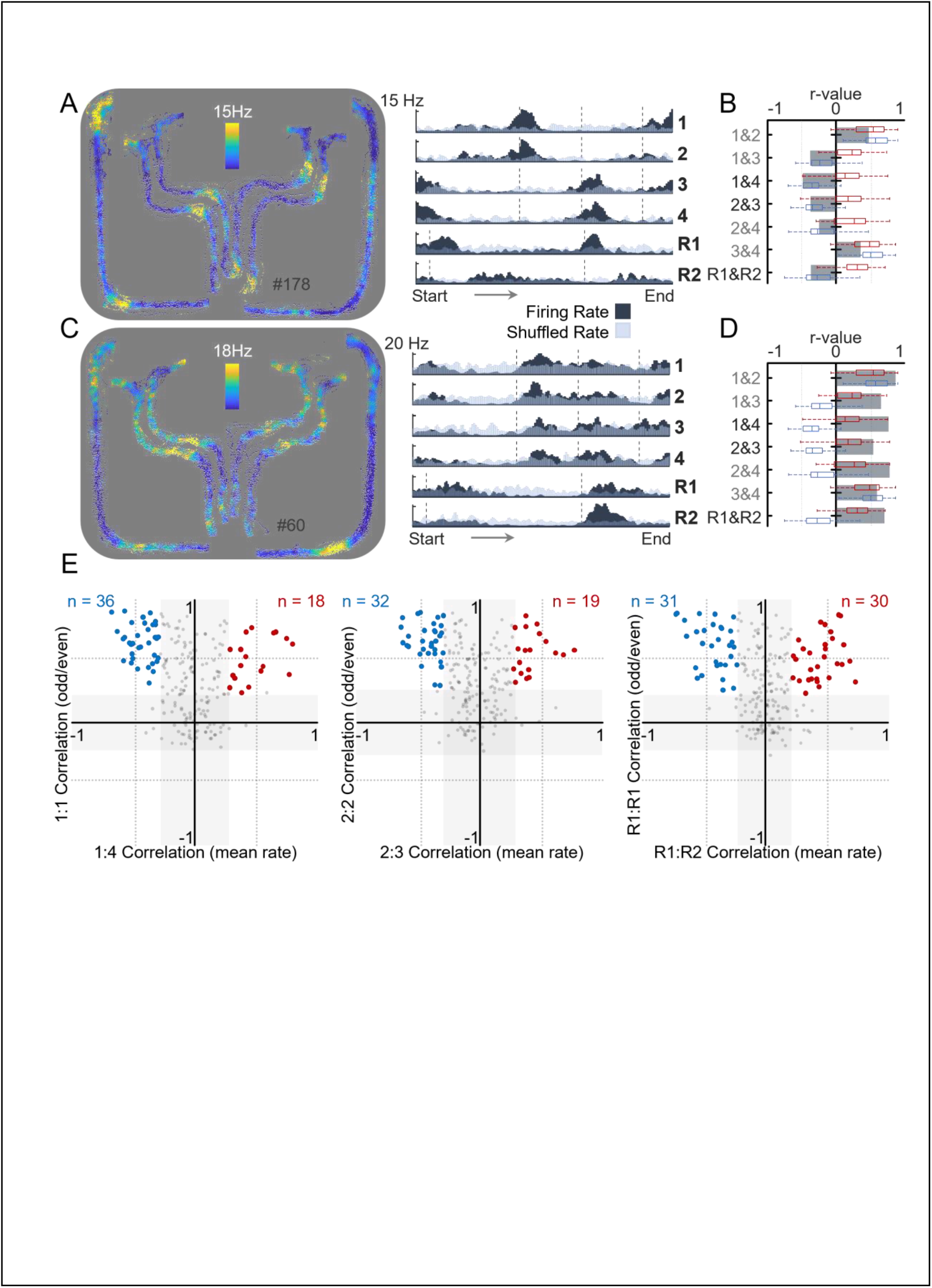
Bifurcation of Firing Rate Correlations Across Dissimilar Path Pairs. A) 2D ratemap of another exemplar neuron exhibiting responses to self-motion (specifically, left-turns). Positional mean firing rate vectors (dark blue) for each route are superimposed with rate vectors generated subsequent to randomized shuffling of spiking data versus tracking data (grey). B) Route-route correlation profiles for the neuron of A are plotted in grey. Superimposed in red box plots are the route-route correlation profiles for all ‘+’ neurons bearing significantly positive correlations for route pairs that demand opposite action sequences (route 1 and 4, 2 and 3, and R1 and R2). Superimposed blue box plots depict correlation profiles for all ‘-‘ neurons bearing significantly negative correlations for route pairs that demand opposite actions sequences. C-D) Same figure layout as in A-B, but for a neuron having positive correlations for route pairs demanding opposite action sequences. E) *Left*: scatterplot of route-route positional rate vector correlations for all neurons for route 1 vs 4 (x-axis) against rate vector correlations computed for odd vs even trials along route 1 . Shaded regions correspond to the mean plus two standard deviations for correlations derived from shuffled data. Neurons labelled in red are significantly positively correlated for route 1 odd-trial versus route 1 even-trial rate vectors and for route 1 versus route 4 rate vectors. Neurons in blue are significantly negatively correlated for route 1 versus route 4 rate vectors, but significantly positively correlated for route 1 odd-trial versus route 1 even-trial rate vectors. *Middle and Right* panels depict the same, but for pairing of route 2 and 3 and pairing of route R1 and route R2. Note that the three sets of paired routes all demand opposite action sequences, such that red or ‘+’ points reflect similarity in rate vectors over routes bearing opposite action sequences while blue or ‘-‘ points reflect opposite patterning in rate vectors over routes demanding opposite action sequences.

A second population of PPC neurons exhibited similar patterns of firing along all four internal and along both return pathways despite their differences in L/R turning series (Figure 2B). Robust firing rate variations for such neurons occurred in similar locations along each route despite the action sequences differing, yielding positive correlations among positional rate vectors. For some neurons with positive path-path correlations, peaks in activity occur along the straight-run portions of path segments, yet do not clearly align to either the linear or angular velocity profiles for those paths.

We defined neurons as being significantly tuned to the structure of the environment based on similarity in firing patterns for path combinations 1 and 4, 2 and 3, and R1 and R2. For each combination, tuning to path structure was defined by inter-route correlation being above or below the mean and 2 standard deviations for an equivalent distribution of correlation values coming from a collection of shuffled data. Further, characterizing neurons as tuned to path structure demanded that they exhibit stable positional rate vectors for odd-numbered versus even-numbered traversals of any single path. Neurons that fell above or below two standard deviations for both criteria were selected for further analyses. For all 3 path combinations, we found neurons that had significantly elevated correlation values for firing rates across same-shaped paths despite differences in the actions taken to move through those paths (referred to as ‘+’ neurons). In contrast, we also found neurons with negative correlations for the same path pairings (referred to as ‘-’ neurons) consistent with tuning to L or R turning behavior.

### PPC Tuning to Self-Motion Explains Differences, but Fails to Explain Similarities in PPC Neuron Activity Across Routes

Given that angular velocities across the path combinations 1/4, 2/3, and R1/R2 are negatively correlated, we considered the possibility that neurons with negative positional firing rate correlations for the same path combinations might exhibit strong tuning to angular velocity. Neurons with positive (‘+’) correlations could reflect tuning to the similarities in shape between these same path combinations. Alternatively, positive correlations might reflect tuning to linear velocities across path locations given that linear velocity was positively correlated across all path combinations.

To examine these potential explanations for the presence of both ‘+’ and ‘-‘ neurons, a generalized linear model approach was adopted (Fig 4A). Here, positional firing rate vectors were modeled using linear and angular velocity as ‘predictors’ in a complete model (cGLM). The normalized mean square error (NMSE) for the cGLM was compared with the 2 ‘partial’ models (pGLM) created by using only linear velocity or only angular velocity as predictors. Tuning to linear velocity was quantified as the difference between the NMSE for the angular velocity pGLM versus the cGLM while tuning to angular velocity was assessed in the same way using the difference between the cGLM NMSE versus the linear velocity pGLM.

**Figure 4.**
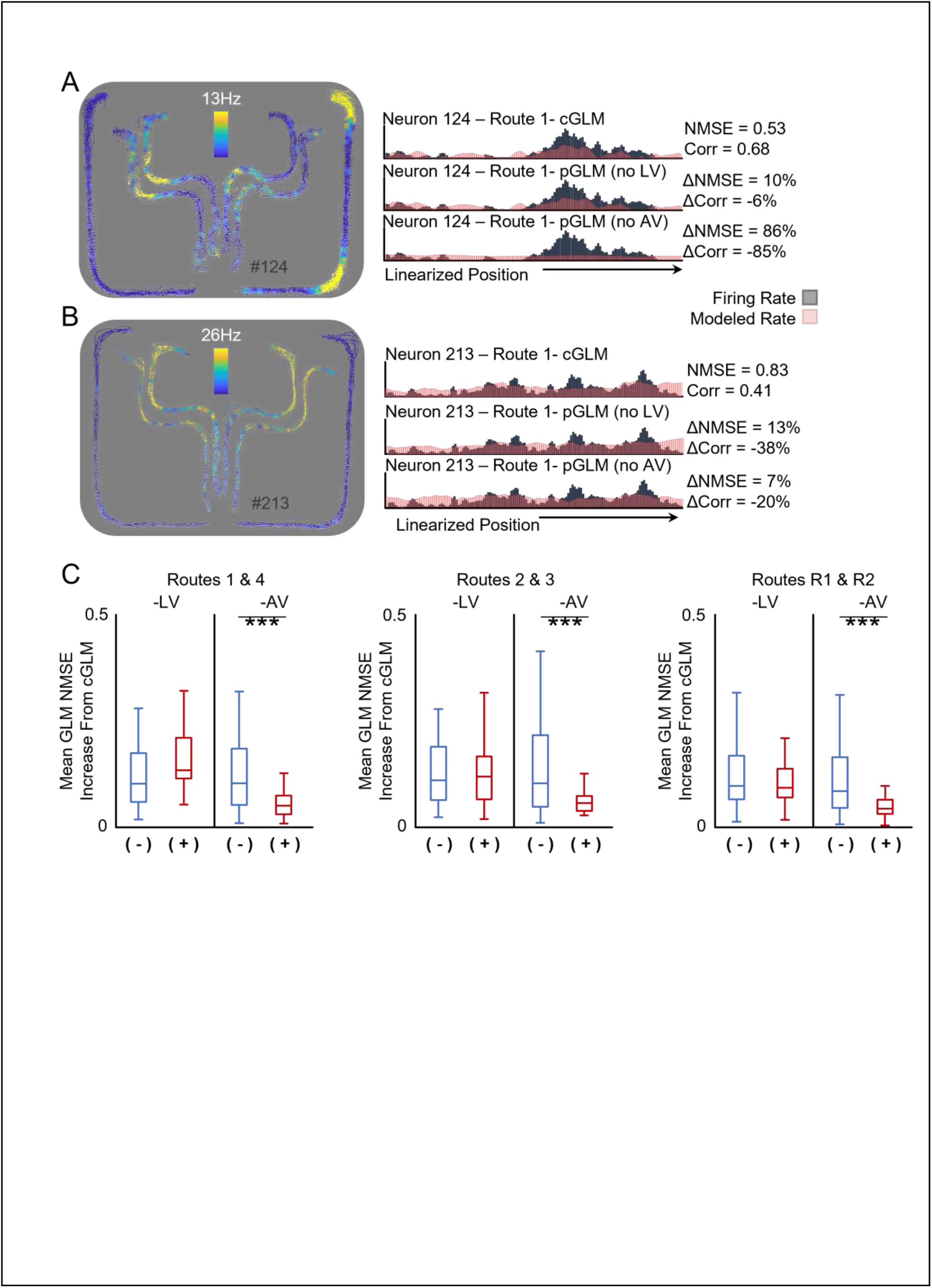
Linear Velocity Fails to Explain Similarity in Firing Rates Across Dissimilar Path Pairs. A) Ratemap for an example ‘-‘ neuron whose firing responses are strongest at R turns. To the right is actual mean firing rate along route 1 (grey) along with the modelled firing rate (red) for the cGLM (top graph) constructed using linear (LV) and angular velocity (AV) as predictors (NMSE = 0.53, r = 0.68). The middle graph depicts the pGLM for the same neuron with the model absent LV. Elimination of LV yields a 10% increase in NMSE with 6% reduction in correlation between the model and actual rate vectors. The bottom graph depicts the pGLM absent AV. Elimination of AV from the model yields a much larger, 86% increase in NMSE with 85% reduction in correlation of actual and model rate vectors. B) Ratemap for an example ‘+‘ neuron with similar patterning of rate across all four internal paths despite differences in their turn sequences. To the right is the actual mean firing rate along route 1 (grey) along with the modelled firing rate (red) for the cGLM (top, NMSE = 0.83, r = 0.41 for correlation of the actual and modelled firing rate vectors. In the middle graph, the pGLM absent LV results in 13% increase in NMSE with 38% reduction in correlation of the actual and modelled rate vectors. The pGLM absent AV (bottom) yields a 7% increase in NMSE with a 20% reduction in correlation. C) Cumulative change in NMSE was calculated across all routes for each neuron’s GLM analysis. The population of neurons above, labelled (+), and below, labelled (-), were tested against each other in a 2-tailed t-test to compare relative contributions of each self-motion variable. Data for routes 1 and 4 (left), routes 2 and 3 (middle), and routes R1 and R2 were combined. For all three route pairings, no significant difference between the ‘+’ and ‘-‘ cell types is observed when LV is eliminated from the model. For all three route pairings, removal of AV from the model increases NMSE significantly more for ‘-‘ neurons than for ‘+’ neurons.

For each of the path combinations, figure 4B depicts the degree to which removal of linear or angular velocity impacted model fitness in pGLMs for the previously defined ‘+’ and ‘-‘ neuron populations. This revealed that ‘+’ and ‘-’ neurons consistently differed only with respect to the strength of angular velocity as a model predictor with ‘+’ neurons exhibiting significantly lower sensitivity (t-test, ***p<0.001). No significant differences were observed for the ‘+’ and ‘-‘ populations in their sensitivity to linear velocity. Thus, the negative correlations in positional firing rates for ‘-‘ neurons may well be explained by their correlation to angular velocity (L/R turning actions) while the positive correlations for ‘+’ neurons cannot be attributed to linear velocity correlations.

## Discussion

In this work, rats displayed excellent spatial working memory performance, dynamically choosing from among multiple, interconnected pathways of a complex path network in order to satisfy task demands. Such behavior set the context for examining the role of PPC in encoding both route shape, and navigational actions. We compared the firing of PPC neurons according to locomotor actions, as quantified by calculation of linear and angular velocities, versus position along routes that were identical in structure, or shape, but differed in the series of turns they demanded. Our analyses focused on three pairs of paths that provided the opportunity to dissociate path shapes from the series of actions required to traverse them. As expected, based on prior work (e.g., McNaughton et al., 1994; Nitz, 2006; Whitlock et al., 2012), our analyses revealed a population of PPC neurons with activity strongly correlated with L versus R turning behavior that exhibit negative correlations of their positional rate vectors across the specific path combinations chosen for analysis. Conversely, we also found a population of PPC neurons with positively correlated positional rate vectors for pairs of task-defined pathways that shared the same shape but were associated with opposite variations in angular velocity during their traversal.

As assessed by a GLM approach using self-motion information (linear and angular velocity) to model positional firing rates along routes, PPC neurons with high negative correlations for path combinations demanding opposite action series appear to have their activity patterns influenced heavily by angular velocity. A separate population of PPC neurons with high positive correlations of their positional rate vectors, when analyzed the same way, did not appear to have the models of their activity patterns impacted by either linear or angular velocity significantly. The latter suggests that while some neurons could be influenced by similar linear velocity series observed across multiple routes, tuning to linear velocity cannot explain the difference between neurons with positive versus negative correlations in their positional rate vectors for different paths. Positively correlated neurons were not unique in the extent that linear velocity could explain their activity. Overall, the results of the GLM analysis find a possible explanation for negative path-path correlations as secondary to angular velocity, or ‘action’, correlations. In contrast, correlation of PPC neurons to self-motion variables fails to explain the recurrence of PPC neuron positional firing rate patterns that generalize across same-shaped paths.

In prior work, PPC has often been considered to function as an ‘action-map’ wherein L and R turning behavior is encoded in the activity of individual neurons, influencing the production of navigational actions through projections to primary and secondary motor cortices (McNaughton et al., 1994; Nitz, 2006; Nitz, 2009; Whitlock et al., 2012; Akrami et al., 2018; Olson et al., 2020; Alexander and Nitz 2023). In the present work, neurons with negative positional rate vector correlations for identically-shaped paths demanding opposite L/R turn series composed a significant proportion of all PPC neurons, supporting models in which PPC functions to generate navigational action plans based on visual or auditory stimuli (Scott et al., 2017) or environmental locations (McNaughton et al., 1994; Whitlock et al., 2012; Olson et al., 2020).

PPC firing correlates to self-motion failed to explain the generalization of complex positional rate vectors across routes demanding different action series. Yet, published work has also demonstrated that PPC neurons can reliably encode locations along a route of the same shape that recurs in different places in an environment, and which demands movement through different headings, environmental locations, and visual landscapes (Nitz, 2006; Nitz, 2009). Such ‘route-centered’ tuning is often complex in its patterning and critical to the interpretation of the present data, robustly discriminates different route locations that share the same navigational action (Nitz, 2009; Nitz, 2012). This is consistent with reporting of context-dependence in PPC neurons that exhibit tuning to self-motion (McNaughton et al., 1994; Whitlock et al., 2012; Harvey et al., 2012).

The present work supports and extends these findings to argue that PPC ‘route-centered’ firing profiles can generalize across different paths that are identical in shape, but which form mirror-images of each other in the horizontal plane of the maze environment and which demand opposite series of L/R turning actions. In this way, one sub-population of PPC neurons can be thought of as encoding the meta-structure, or topology, shared by routes. It follows that repeats in PPC ensemble activity patterns will be distributed in non-random fashion within path networks, such as city grids, that afford the same-shaped routes from many environmental locations (e.g., all-L-turn or all-R-turn trips “around the block” for all blocks of a city grid). Based on this, we suggest that such recurrence in PPC ensemble patterns, in combination with hippocampal ensemble patterns discriminating all environmental locations, can form the basis for learning the structural layout of an environment’s path network structure. Our results evidence the co-existence of PPC neuron populations with tuning to actions and PPC neuron populations with tuning to much more abstract spatial series that recur during navigation in environments with non-random path-network structure which we refer to here as the ‘meta-structure’ of that environment.

The encoding of meta-structure could be expected to be found in the brain given the large-scale understanding of complex space such as that influencing route choice when navigating cities of different layouts (Sevtsuk and Basu, 2022). Experience with diverse meta-structures may explain then, why navigation ability is greater in individuals who previously had learned city-layouts of high entropy (Coutrot et al., 2022). Furthermore, the results are in accordance with human lesion studies examining ‘topographic amnesia’ in which patients with PPC damage are unable to both learn new routes and how they relate to the spaces they connect to (De Renzi et al. 1977).

---

**Supplemental Figure 1.**
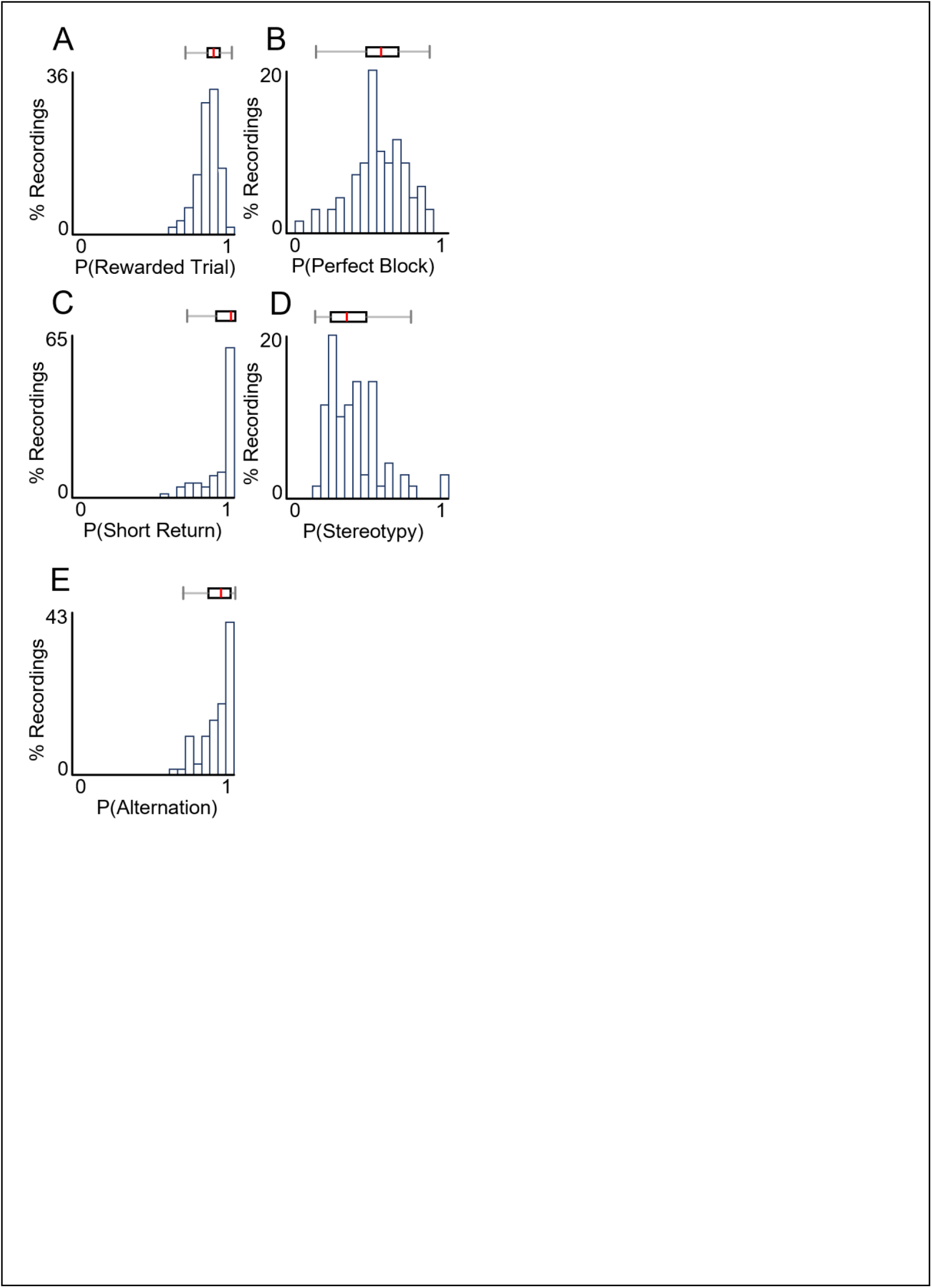
Details of Animal Performance on Working Memory Task. Compilation of performance on the find-all-4 condition. A) Probability of getting a reward mean = 0.848 s.d = 0.0688. B) Probability of completing a block without error mean = 0.566 s.d. = 0.1679. C) Probability of choosing the shorter of two return arms mean = 0.9197 s.d. = 0.1075. D) Probability of perfect blocks having stereotypy in the sequence of routes utilized. Here the probability of the maximally observed route sequence in each recording is used to measure the degree of stereotypy. mean = 0.4020 s.d. = 0.1730. E) Probability of alternating at the first decision point on subsequent trials. mean = 0.8973 s.d. = 0.0916

**Supplemental Figure 2.**
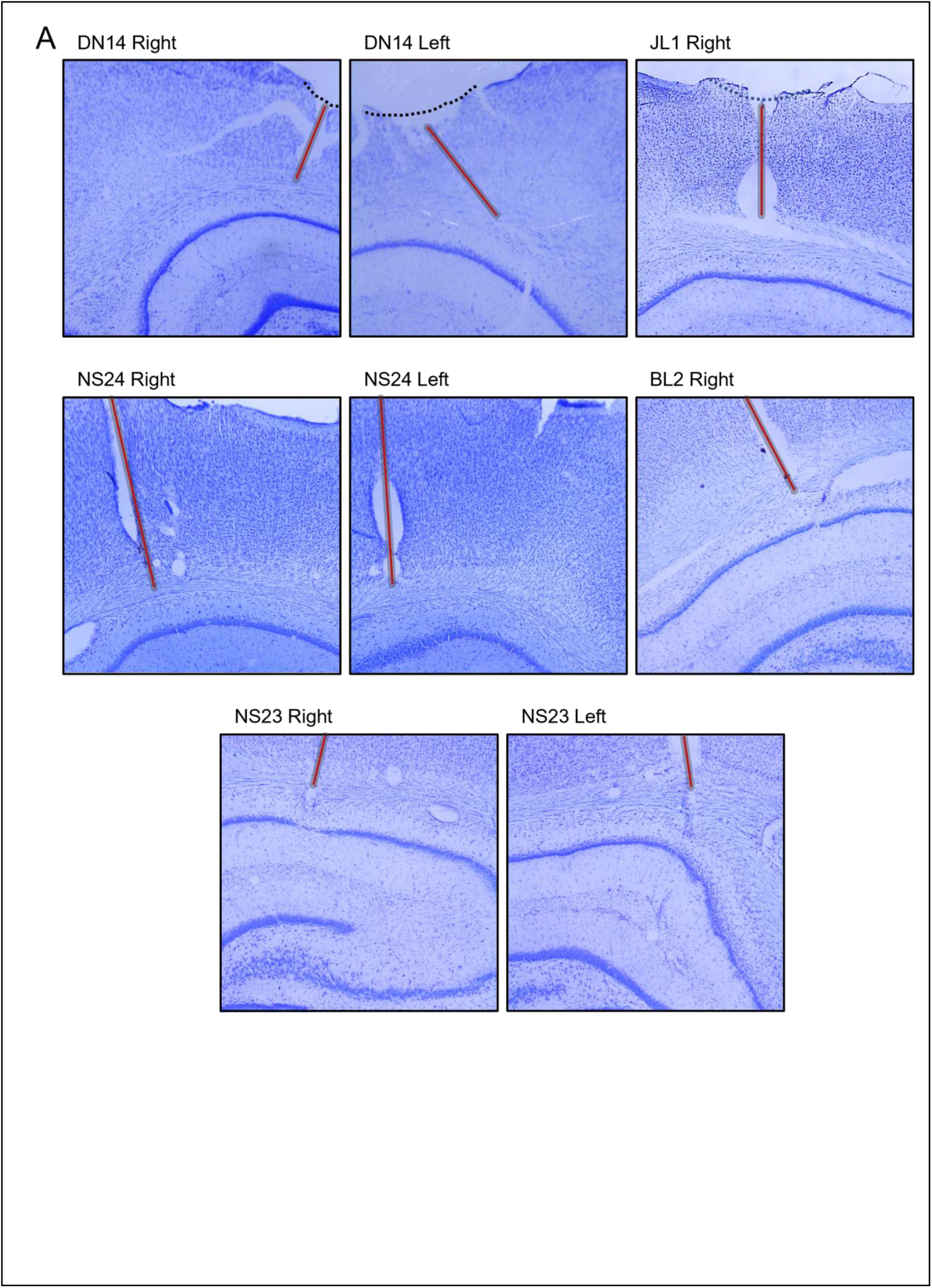
Summary of Recording Site Histological Data. Nissl stained brain sections from each animal depicting their tetrode bundle trajectories. Anatomical locations for recordings sessions included in the dataset are highlighted in red.

**Supplemental Figure 3.**
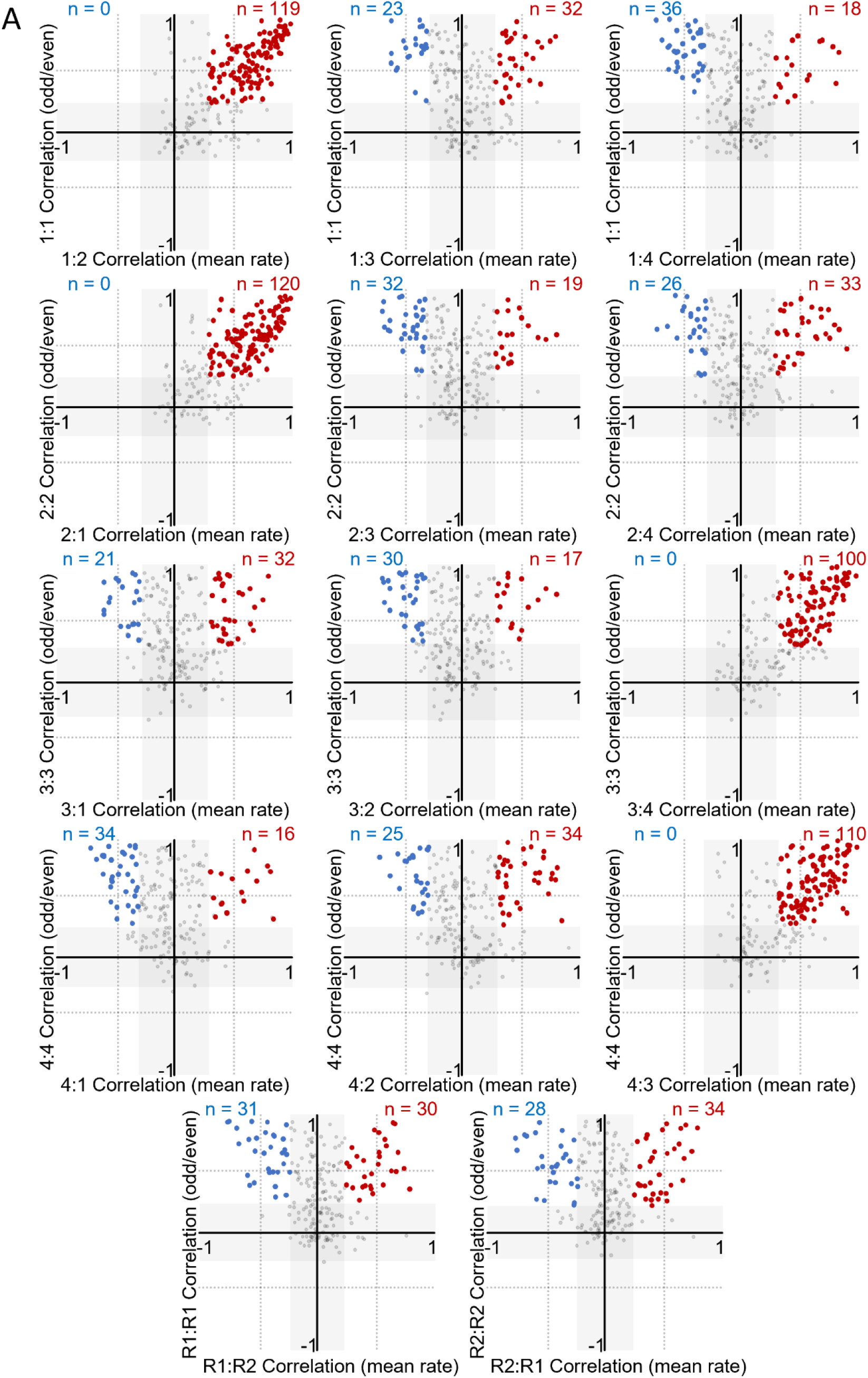
Scatterplots for correlation values of individual neuron’s positional rate vectors for all route combinations (x-axis) against within-route positional rate vector correlations for odd versus even trials (y-axis, note that route 1/4, route 2/3, and route R1/R2 data are repeated from figure 3E). Shaded regions in each correspond to the mean plus two standard deviations for correlations derived from shuffled data (randomized circular shuffling of rate vectors). Neurons labelled in red are significantly positively correlated for both the within-path and cross path rate vector correlations. Neurons in blue are significantly negatively correlated for cross-route correlations and significantly positive for within-route correlations. Neuron counts for each grouping (above or below) are presented above the route-route projections.

